# Investigating the dynamics of precursor availability for sesquiterpenoid production in the cytoplasm and plastid of *Chlamydomonas reinhardtii*

**DOI:** 10.1101/2025.06.15.659761

**Authors:** Malak N. Abdallah, Gordon B. Wellman, Sebastian Overmans, Kyle J. Lauersen

## Abstract

The green alga *Chlamydomonas reinhardtii* is an interesting organism for production of heterologous isoprenoid natural products through metabolic engineering as it can grow photosynthetically and represents a plant-like cell environment for synthase expression. Here, we investigated expression of the patchoulol synthase from *Pogostemon cablin* (Benth) (*Pc*PS) to probe the production of sesquiterpenoids from different sub-cellular compartments of the algal cell. We observed that knock-down of squalene synthase (SQS) in the cytoplasm did not affect sesquiterpene production titers in the plastid, suggesting no backward movement of FPP from cytoplasmic pools. Fusion of *Pc*PS with a farnesyl pyrophosphate synthase (FPPS) did not enhance patchoulol production in the cytoplasm but did in the plastid, as also reported elsewhere. Secondary transformations with cytoplasmic and chloroplast-localized *Pc*PS constructs increased synthase titers and improved patchoulol yields from both compartments. In multi-parallel photobioreactor experiments with patchoulol extraction through two-phase solvent contact, different daily patchoulol production behaviors were observed based on subcellular localization of the *Pc*PS and carbon source. Those with plastid localization of *Pc*PS-FPPS exhibited continued production into later stages of cultivation, while cytoplasm-localized *Pc*PS with SQS k.d. had the highest production rates in mid-logarithmic phase with low productivity in later growth stages. Cultivations in high-density photobioreactors with nutrient-enriched medium generated 15 mg patchoulol/L culture from CO_2_, the highest yield reported from the alga to date. The results indicate that the algal cell has flexible pools of isoprenoid precursors in different subcellular compartments from which the biotechnologist can design an expression strategy tailored to carbon-feeding regimes.

## 1. Introduction

Isoprenoids, also known as terpenes or terpenoids, are a structurally diverse class of naturally occurring hydrocarbon molecules (Kirby & Keasling, 2009; Lauersen, 2019). These compounds play crucial roles in cellular processes, including photosynthesis, and are the precursors of carotenoids, sterols, and chemicals like ubiquinone (Boncan et al., 2020). All Isoprenoids are generated naturally from the condensation of the universal five-carbon (C5) precursors isopentenyl pyrophosphate (IPP) and its isomer dimethylallyl pyrophosphate (DMAPP) into the higher carbon chain precursors: geranyl pyrophosphate (GPP, C10), farnesyl pyrophosphate (FPP, C15), and geranylgeranyl pyrophosphate (GGPP, C20) (Zhao et al., 2013; Wichmann et al., 2018). Through the action of enzymes known as terpene synthases, these precursors are converted into the myriad of different terpenes found in nature. Terpenes are classified according to the number of carbons: hemiterpenes (C5), monoterpenes (C10), sesquiterpenes (C15), diterpenes (C20), triterpenes (C30), and tetraterpenes (C40) (Lohr et al., 2012; Lauersen et al., 2016; Wichmann et al., 2018). Isoprenoids have a range of human-use applications, from pharmaceuticals and nutrition (Kirby & Keasling, 2009) to biofuels (Kang et al., 2022). However, cost-effective and sustainable large-scale extraction of terpenoids from natural sources remains challenging (Wichmann et al., 2020). Many terpenes are plant-derived, occur in low quantities, or are difficult to extract and purify from their native hosts (Boncan et al., 2020; Gangl et al., 2015). Bioengineering of microbes as alternative production vehicles for isoprenoids has developed into a mature technology (Kirby & Keasling, 2009; Gangl et al., 2015; Leavell et al., 2016; Gutiérrez et al., 2024). Terpene synthases can be overexpressed in microbial hosts, wherein they are able to convert cellular precursors into a desired chemical identical to that in their native host. The precursor availability can also be further enhanced through metabolic engineering.

Bacteria (Davies et al., 2014) and yeast (Kirby et al., 2014; Qin et al., 2024) are widely used hosts for isoprenoid production due to their robust genetic tractability and ease of cultivation on an industrial scale. However, these microorganisms rely on an organic carbon source, such as glucose, which necessitates the geographical positioning of fermentation facilities near agricultural production sources of sugars (Davies et al., 2014). Therefore, commercial production of isoprenoids from fermentation may not be aligned with resource management goals in certain geographies (Lauersen, 2019). Photosynthetic microbes, as an alternative, have the potential to convert CO_2_ into their biomass and coincident bioproducts. As CO_2_ is an abundant and readily available waste stream, engineered algae that produce terpenes pose the possibility of waste-revalorization into these specialty chemicals concomitant with algal biomass production processes (Chu, 2012).

The eukaryotic microalga *C. reinhardtii* has emerged as a new host organism for metabolic engineering over the past decade. As a photosynthetic cell, it natively accumulates carotenoid pigments as terminal isoprenoids, making it primed as a plant-like cell for precursor supply to heterologous terpenes. *C. reinhardtii* has been used for the heterologous production of hemi-, mono-, sesqui-, and diterpenes (Lauersen et al., 2016, 2018; Wichmann et al., 2018; Abdallah et al., 2022; Kang et al., 2022; Wichmann et al., 2022; Yahya et al., 2023; Gutiérrez et al., 2024)*. C. reinhardtii* produces all cellular isoprenoids from its native 2-C-methyl-D-erythritol 4-phosphate (MEP) pathway, which is encoded in the nuclear genome and enzymes localized to its plastid (Lohr et al., 2012). Improvements in *C. reinhardtii* metabolic engineering have been achieved through genetic optimization strategies, including overexpression of rate-limiting genes in the MEP pathway (Lauersen et al., 2018), inhibiting genes that compete for PP precursors (Lauersen et al., 2016; Abdallah et al., 2022; Einhaus et al., 2022), and over-expression of terpene synthases (Lauersen et al., 2016).

In addition to the requirement of high sesquiterpene synthase overexpression to mediate production, the challenge of generating a substantial precursor pool of FPP limits the biosynthesis of most sesquiterpenoids (Lei et al., 2021). Studies of sesquiterpene production have shown that the C15 farnesyl pyrophosphate (FPP) is present and accessible for metabolic engineering in the cytoplasm; however, other terpene precursors are not present or accessible (Wichmann et al., 2018). In the plastid, however, IPP, DMAPP, GPP, and GGPP are available and accessible for metabolic engineering. Only trace amounts of FPP were found to be accessible for heterologous product generation in the algal plastid. Co-expression of an FPP synthase from yeast (*Sc*ERG20) and the *Abies grandis* bisabolene synthase (*Ag*BS) in the chloroplast was reported to facilitate bisabolene production by increasing the pool of available FPP in this compartment (Wichmann et al., 2022), as well as for many other sesquiterpenoids (Gutiérrez et al., 2025).

In the cytoplasm, it has been shown that knockdown of the native squalene synthase (SQS k.d.) enhances heterologous sesquiterpenoid production as SQS is the main competitor for FPP in this location (Abdallah et al., 2022; Gutiérrez et al., 2024; Wichmann et al., 2018). Leveraging different organelles for terpene production can potentially reduce metabolic interference and improve precursor availability towards increasing the yield of isoprenoids (Jia et al., 2020; Chen et al., 2023).

This work explores the subcellular dynamics of sesquiterpene production in *C. reinhardtii* by investigating the cross-talk of metabolic precursor pools between the cytoplasm and chloroplast to better understand their metabolic dynamics. The patchoulol synthase from the plant *Pogostemon cablin* (*Pc*PS) (Deguerry et al., 2006) was used as a sesquiterpenoid synthase to explore the cellular capacity for heterologous sesquiterpenoid production in either compartment. Patchoulol production dynamics were examined in strains with *Pc*PS overexpressed and localized in cytoplasm or plastid, with and without knockdown of the native cytoplasmic squalene synthase (SQS) to investigate any effect this may have in the plastid. The patchoulol production dynamics were further investigated in two strains where different engineering strategies led to similar total patchoulol titers: overexpression of the *Pc*PS in the algal cytoplasm with SQS k.d. or *Pc*PS-FPPS fusions localized in the plastid. These strains were subjected to photobioreactor cultivations with CO_2_, acetic acid, or both as carbon sources, and the dynamics of patchoulol production were demonstrated through daily sampling. The results indicate that the production dynamics of heterologous sesquiterpenes depend on the subcellular localization of synthases and the dynamic nature of their precursors in response to carbon sources.

## 2. Materials and Methods

### 2.1. Algal culture condition

The *C. reinhardtii* strain UPN22 was used in the experiments described here. UPN22 is a derivative strain of UVM4 (Neupert et al., 2009) that has been previously engineered to use phosphite as the sole source of phosphorous and nitrate as a nitrogen source (Abdallah et al., 2022). ‘UPN22 SQS k.d.’, as referred to throughout this work, is a derivative strain, which contains an artificial micro RNA (amiRNA) for knockdown of the *C. reinhardtii* squalene synthase (Uniprot ID: A8IE29) as previously reported using the pOpt2_cCA-gLuc_i3-SQS_Spect plasmid (Wichmann et al., 2018; Abdallah et al., 2022; Gutiérrez et al., 2024). Microalgal cultures were routinely maintained on solidified agar TAPhi-NO_3_ medium (Abdallah et al., 2022) at 150 µmol photons m^−2^ s^−1^ light intensity or in liquid TAPhi-NO_3_ medium with 120–190 rpm agitation in shake flasks or microtiter plates. The growth of algae was analyzed by flow cytometry, as described below.

### 2.2. Genetic tools, design, synthesis, and screening

The plasmids used in this work are listed in Supplementary Table 1, and their full annotated sequences can be found in Supplemental File 1. All cloning and plasmid linearization were performed with FastDigest (Thermofisher) restriction enzymes, and Quick Ligase (New England Biolabs), following the manufacturer’s protocols. Plasmids were maintained in chemically competent *Escherichia coli* DH5α transformed by heat shock. The *Pogostemon cablin* patchoulol synthase (*Pc*PS, UniProt Q49SP3) plasmids for the algal nuclear genome were adapted from Lauersen et al. (2016) but updated to the pOpt3 plasmids (Gutiérrez et al., 2022). Briefly, *Pc*PS was subcloned into *Bam*HI-*Bgl*II of a pOpt3_mTFP1_Ble plasmid. Plastid targeting was achieved with the N-terminal chloroplast-targeting peptide (CTP) derived from the *C. reinhardtii* photosystem I reaction center subunit II (PsaD). Further fusions were made with C-terminal yeast FPP-synthase (*Saccharomyces cerevisiae Sc*ERG20, Uniprot ID: P08524) as previously reported (Wichmann et al., 2022; Gutiérrez et al., 2025). Transgene expression from the algal nuclear genome is assisted by fluorescent reporter fusion proteins that can identify highly expressing transformants on agar plates (Lauersen, 2019; Gutiérrez et al., 2022). In this work, the monomeric teal fluorescent protein 1 (mTFP1) was fused to the *Pc*PS as a reporter, and selection was achieved with the bleomycin resistance cassette (*Sh*Ble); the monomeric yellow fluorescent protein, mVenus, and paromomycin resistance (*APHVIII*) were also used in some plasmids for secondary transformation. The reporters were used to enable plate-level fluorescence detection in transformant colonies (Gutiérrez et al., 2022), as described below.

### 2.3. Transformation and positive transformant screening

Nuclear transformation of all strains was performed by glass bead agitation as previously described (Kindle, 1990). UPN22 and UPN22+SQS k.d. strains were transformed with various *Pc*PS expressing plasmids #1 to #6 (**Supplementary Table 1**). Transformations were plated on selective medium (bleomycin 10 mg L^−1^ and/or spectinomycin 200 mg L^−1^) and left under constant illumination for ∼7 d before colony picking. A PIXL robot (Singer Instruments, UK) was used to pick 384 colonies per transformation and transfer them to TAPhi-NO_3_ agar plates with bleomycin (10 mg L^−1^) and/or spectinomycin (200 mg L^−1^), depending on plasmid selection marker as previously described (Abdallah et al., 2022; Gutiérrez et al., 2022). Afterward, colonies were replicated to new plates by stamping using a ROTOR robot (Singer Instruments, UK), as well as to plates containing 150 mg L^−1^ amido-black for fluorescence screening, as previously described (Abdallah et al., 2022; Gutiérrez et al., 2022). Briefly, chlorophyll fluorescence was captured to normalize colony presence/absence on plates with 475/20 nm excitation and DNA gel emission filter (640/160 nm) for 3 s exposure. The mTFP fluorescence signal was captured with 475/20 nm excitation and 510/10 nm emission filters for 2.5 min (Analytik Jena Chemstudio Plus GelDoc). Transformants with high fluorescence signals were selected and inoculated in liquid culture to quantify patchoulol productivity using dodecane overlays and gas chromatography-mass spectrometry (GC-MS), as described below. The predicted molecular masses of expressed proteins were confirmed by in-gel fluorescence on sodium dodecyl sulfate-polyacrylamide gels (SDS-PAGE), as previously described (Gutiérrez et al., 2022). To enhance heterologous patchoulol production, one selected transformant generated from each Plasmids 03 and 06 was further transformed with the Plasmids 07, 08, or 09, followed by selection on TAPhi-NO_3_ medium with bleomycin 10 mg L^−1^, paromomycin 15 mg L^−1^ and spectinomycin 200 mg L^−1^. Primary positive transformants and colony picking were performed as described above. Chlorophyll and mVenus fluorescence signals were captured, whereby mVenus signals were captured with 504/10 nm excitation and 530/10 nm emission filters for 30 s (Analytik Jena Chemstudio Plus gel doc). Screened transformants with mVenus and mTFP1 signals containing both transgenes of interest were then inoculated in liquid medium, and their patchoulol production quantified.

### 2.4. Quantifying patchoulol production of transformants

After fluorescence screening of the agar plates, UPN22 and UPN22+SQS k.d. transformants expressing *Pc*PS variants were screened for heterologous patchoulol production. Colonies from agar plates were inoculated in 1 mL liquid TAPhi-NO_3_ medium in 24-well microtiter plates and grown for 3 d under constant light (150 µmol m^−2^ s^−1^) and 190 rpm orbital shaking. On day three, 500 µL of each transformant was transferred into triplicate wells of a 6-well microtiter plate, each well containing 4.5 mL TAPhi-NO_3_. 500 µL of n-dodecane overlay was added to each well, acting as a patchoulol accumulating layer. The algae were grown for 6 d on orbital shakers (120 rpm) with 150 µmol m^−2^ s^−1^ illumination. Subsequently, cell concentrations were analyzed by taking a subsample of the culture from each well for flow cytometry, as described below. The remaining content of three triplicate wells was pooled in 15 mL test tubes. The tubes were centrifuged at 1000 x *g* for 5 min to separate dodecane and algal cultures. The resulting dodecane supernatants were transferred into separate 2 mL microcentrifuge tubes and stored at -20°C until further analysis using GC-MS (Lauersen et al., 2018).

### 2.5. Multi-parallel photobioreactor cultivations

Two transformants of UPN22+SQS k.d. strain with dual expression of *Pc*PS from different subcellular compartments (Plasmids # 3 + 8 and, 6 + 9) hereafter referred to as ‘cytoplasm-cytoplasm’ and ‘chloroplast-chloroplast’, were used in this experiment. Each transformant line was tested separately and grown in Algem photobioreactors (Algenuity, UK). The selected transformants were maintained on TAPhi-NO_3_ agar plates and scaled up in liquid culture by sequential transfer into 1 mL, 6 mL, and 50 mL of the same liquid medium and grown in Erlenmeyer flasks under ∼100 μmol photons m^−2^ s^−1^ and 120 rpm shaking before bioreactor cultivation. Before experiments, transformants were cultivated in 400 mL TAPhi-NO_3_ medium shaken at 100 rpm and exposed to 325 μmol photons m^−2^ s^−1^ on a 12h:12h dark: light cycle at 25 °C. In each experiment, one transformant was grown in 2 L Erlenmeyer flasks with a total volume of 400 mL culture shaking (100 rpm) at 23 °C and 325 μmol photons m^−2^ s^−1^ in six different growth conditions for 6 d (Supplementary Table 2). For acetate-based growth, TA2Phi-NO_3_ medium was used; for photoautotrophic cultivation, T2Phi-NO_3_ medium - without acetate but containing 2X phosphite concentration (Lauersen et al., 2016) - supplied with 3% CO_2_ in air. For mixotrophic growth, TA2Phi-NO_3_ + 3% CO_2_. To capture patchoulol produced by the algal cells during growth, 20 mL of the perfluorinated amine FC-3283 (3M, sourced via Acros Organics, Geel, Belgium) was added to each culture as an underlay (Overmans & Lauersen, 2022; De Freitas et al., 2023). Daily, 500 μL of FC-3283 was sampled from each bioreactor flask to quantify the patchoulol product accumulated in the underlay. All FC-3283 samples were immediately stored at -20 °C until the end of the experiment and processed collectively for GC-MS analysis. 1 mL of the algae culture was sampled from each flask to determine algal-cell densities using flow cytometry, as described below.

### 2.6. High-density autotrophic cultivation for patchoulol production

Two transformants, with ‘cytoplasm-cytoplasm’ and ‘chloroplast-chloroplast’ subcellular localized *Pc*PS constructs (Plasmids # 3 + 8 and, 6 + 9) were cultivated in a 6xPhi medium with dodecane overlay in a CellDEG HD100 bioreactors. Starter cultures were grown until the mid-log phase in TAPhi-NO_3_ media, centrifuged at 1000 x *g* for 4 min, and the supernatant discarded. Cells were resuspended in a 6xPhi medium and diluted to ∼5x10^5^ cells mL^−1^. The dilute culture was divided between three independent HD100 cultivators, containing 200 mL of dilute culture and 20 mL dodecane added to form an overlay. Cultivators were placed onto the CellDEG platform and gently shaken at 90 rpm to avoid contact between the dodecane overlay and the gas exchange membrane. Cultures were grown with light and CO_2_ profiles (see Supplementary Table 3) and maintained at ∼20–22 °C during the experiment. Balanced white light was used, and intensities were measured at the approximate position of the culture surface with a C-7000 handheld spectrometer (Sekonic, USA). 100 µL of culture was sampled in triplicate every 2 d for cell counts by flow cytometry, as described below. One replicate of each strain was harvested on day 7, and the dodecane overlay was separated for GC-FID quantification of patchoulol, as described below. The remaining two replicates were maintained at higher light intensity for an additional 7 d for maximum patchoulol production before harvesting for GC-FID quantification.

### 2.7. Algal cell density estimation by flow cytometry

Algal growth was analyzed by flow cytometry using an Invitrogen Attune NxT flow cytometer equipped with a Cytkick microtiter plate autosampler unit (Thermo Fisher Scientific, UK), as previously described (Overmans & Lauersen, 2022). Briefly, 10 µL from each biological duplicate (*n*=2) was diluted 1:100 with 990 µL of 0.9% NaCl solution. 200 µL of each diluted sample was loaded into three separate wells of a 96-well microtiter plate and measured in technical triplicates (n=3). The plate was loaded into the autosampler, and the samples were mixed three times immediately before analysis. The first 25 µL of each sample was discarded to ensure a stable flow rate. Data acquisition automatically stopped when 50 μL from each well was analyzed. Population clustering was performed using Attune NxT software v3.2.1 (Life Technologies, USA).

### 2.8. Gas Chromatography-Mass Spectrometry

Patchoulol was quantified by transferring 90 µL of each collected solvent sample into triplicate 2 mL GC vials, dodecane overlay for plate-based screening and CellDeg experiments, or FC-3283 underlay for Algem photobioreactor experiments. A patchoulol standard (Cayman Chemical Company, USA, catalog# 18450) calibration curve in dodecane or FC was used for linear-range quantification of patchoulol in the range 10–200 mg/L. 250 mg/L of cedrene (Sigma-Aldrich, USA catalog #22133) was added as an internal standard to each sample and patchoulol standard. All samples were analyzed using an Agilent 7890A gas chromatograph (GC) equipped with a DB-5MS column (Agilent J&W, USA) attached to a 5975C mass spectrometer (MS) with a triple-axis detector (Agilent Technologies, USA). A previously described GC oven temperature protocol was used (Overmans & Lauersen, 2022). All GC-MS measurements were performed in triplicate, and chromatograms were manually reviewed for quality control. Gas chromatograms were evaluated with MassHunter Workstation software version B.08.00 (Agilent Technologies, USA).

## 3. Results & Discussion

### 3.1. Sesquiterpenoid production in different subcellular compartments

The patchoulol synthase *Pc*PS was used, with and without a plastid-targeting peptide, to investigate the cellular capacity to produce heterologous sesquiterpenoids in cytoplasm or chloroplast **(Figure 1A)**. These were performed in a parental ‘UPN22’ as well as the ‘UPN22+SQS k.d.’ strains to investigate what effect SQS k.d. had on plastid localized sesquiterpenoid production **(Figure 1A-C)**. In addition, variants of *Pc*PS with fusion to a yeast FPPS (*Sc*ERG20) were also expressed in both strains and localized to both subcellular compartments **(Figure 1A-C)**. The FPPS has been shown to increase FPP pools if IPP and DMAPP are available, which will result in higher patchoulol production (Wichmann et al., 2022) (Plasmids are listed in **Supplementary Table 1)**.

**Figure 1.**
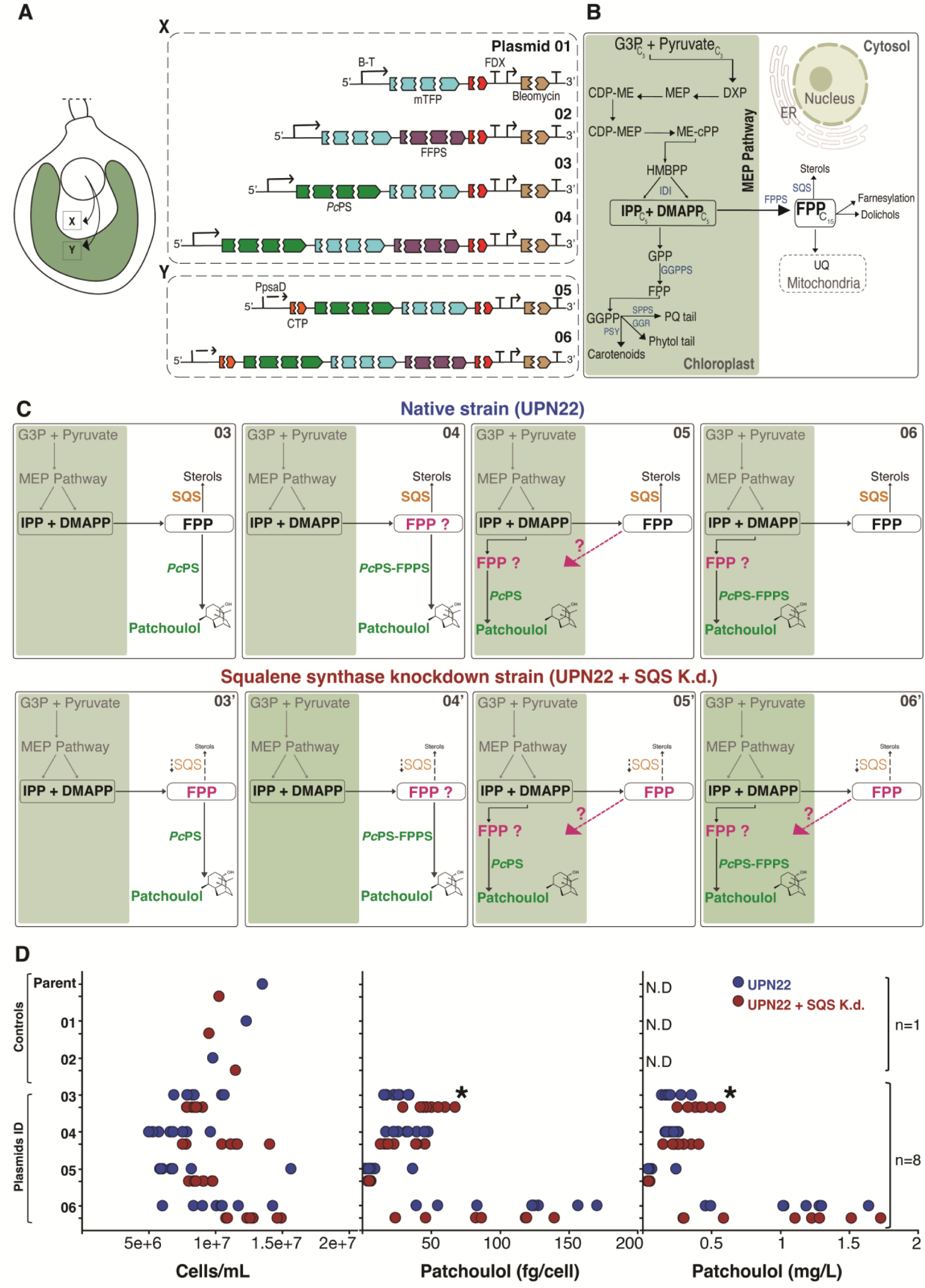
Cytoplasm and chloroplast localized *Pc*PS fusion vectors and their respective patchoulol productivity. **A.** Sketch of a *C. reinhardtii* cell with nucleus (circle), and plastid (green) indicating that nuclear transformed constructs target the patchoulol synthase (*Pc*PS) to either cytoplasm (X) or plastid (Y) with diagrams of plasmids 01-06 used to transform either UPN22 (native state) or UPN22+SQS k.d. strains. PcPS was fused to mTFP fluorescent reporter alone or the yeast FPP synthase (*Sc*ERG20) and localized to the algal cytoplasm (X) or targeted to the plastid (X). All plasmid sequences and their annotations are presented in Supplemental File 1. **B.** Diagram of the native algal non-mevalonate (MEP) pathway, abbreviations below. **C.** Modifications of MEP pathway and the engineered production of patchoulol in the native strain UPN22 and squalene synthase knockdown strain UPN22+SQS k.d. transformed with various plasmids as indicated. **D.** Patchoulol productivity of cytoplasmic and chloroplast targeted constructs in either UPN22 (red) or UPN22+SQS k.d (blue). Individual transformants (n=8) were cultivated with dodecane overlay in triplicate (n=3) and after six days of cultivation patchoulol productivities determined by GCMS. N.D. indicates patchoulol not detected. Glycerol-3-phosphate (G3P), 1-deoxy-D-xylulose 5-phosphate (DXP), 2-C-methyl-D-erythriol 4-phosphate (MEP), 4-diphosphocytidyl-2-C-methylerythritol (CDP-ME) and -2-phosphate (CDP-MEP), 2-C-methyl-D-erythritol 2,4-cyclodiphosphate (MecPP), 4-Hydroxy-3-methyl-but-2-enyl diphosphate (HMB-PP), isopentenyl diphosphate (IPP)/dimethylallyl diphosphate (DMAPP) isomerase (IDI), geranyl pyrophosphate (GPP), geranylgeranyl pyrophosphate (C20) and its synthase (GGPPS), geranylgeranyl reductase (GGR), farnesyl diphosphate (FPP, C15) and its synthase (FPPS), solanesyl diphosphate (SPP, C45) and synthase (SPPS), phytoene synthase (PSY), plastoquinone (PQ), phytoene synthase (PSY), squalene synthase (C15), Ubiquinone (UQ), endoplasmic reticulum (ER). The single significant difference is noted by an asterisk (*p* = 0.0002).

As expected, the empty vector (Plasmid 01) and FPPS alone (Plasmid 02) did not result in detectible patchoulol accumulation **(Figure 1D)**. Transformation with Plasmid 03 yielded UPN22 strains with an average of 24 fg patchoulol /cell and UPN22+SQS k.d. strains with 50 fg patchoulol/cell **(Figure 1D)**. It was observed that the highest UPN22 transformant generated up to 34 fg patchoulol/cell, but this was significantly greater in UPN22+SQS k.d. with a maximum titer of 68 fg patchoulol/cell **(Figure 1D**, **t(DF) = 12.949, *p* = 0.0002). The increased patchoulol titer mediated by SQS k.d. is consistent with previous findings and expected as it reduces competition for the FPP substrate** (Wichmann et al., 2018; Abdallah et al., 2022; Gutiérrez et al., 2024). The *Pc*PS construct with FPP-synthase (*Sc*ERG20) fusion localized in the cytoplasm (Plasmid #04) resulted in an average patchoulol production of 33 and 27 fg patchoulol /cell in UPN22 or UPN22+SQS k.d., respectively **(Figure 1D)**. This is in line with previous findings that the FPPS does not improve cytoplasmic pools of FPP, likely as IPP and DMAPP are not exported from the plastid (Lauersen et al., 2016; Wichmann et al., 2022; Gutiérrez et al., 2025). The PsaD chloroplast targeting peptide (CTP) was used on Plasmids 05 (*Pc*PS only) and 06 (*Pc*PS with FPPS fusion) to test the ability to produce patchoulol in the chloroplast **(Figure 1C 05+06)**. Only negligible amounts of patchoulol were produced from Plasmid 05 in both UPN22 and UPN22+SQS k.d. strains: 10 and 5 fg patchoulol/cell, respectively **(Figure 1D).** The FPPS-fusion, however, produced an average of 110 and 80 fg patchoulol /cell in UPN22 and UPN22+SQS k.d., respectively **(Figure 1D).** The highest-producing line yielded 170 and 139.5 fg patchoulol/cell for UPN22 and UPN22+SQS k.d., respectively **(Figure 1D).** These findings are in agreement with results reported for bisabolene and other sesquiterpenoids (Wichmann et al., 2022; Gutiérrez et al., 2025). Co-overexpression of an FPPS and the patchoulol synthase in tobacco plastids also improved patchoulol production several-fold (Wu et al., 2006). Importantly, we were interested in determining if SQS k.d. led to feedback on the plastid and the variance in FPP availability there. Based on the patchoulol titers quantified here, no difference in patchoulol yields was observed when *Pc*PS was plastid localized in the SQS k.d. strain compared to the parental, suggesting this modification does not impact plastid FPP pools **(Figure 1C** 05 compared to 05’ and **D**).

### 3.2. Loading of synthase titers in each compartment

We sought to determine if different cellular compartments could be simultaneously used for heterologous patchoulol production and whether improved rates could be achieved through secondary transformation and further synthase expression **(Figure 2A).** Selected transformants of Plasmids 03 and 06 in UPN22+SQS k.d. were further engineered by subsequent transformation with cytoplasmic or chloroplast-localized *Pc*PS constructs. These were fused to mVenus for separate screening from the original mTFP1-fused versions **(Figure 2A)**. This strategy allowed the generation of transformants with *Pc*PS entirely in the cytoplasm, in the cytoplasm and plastid, or entirely in the plastid **(Figure 2)**. After confirming correct target transgene expression **(Figure 2B),** selected transformants of each combination (n=8) were grown in liquid culture with dodecane overlay for patchoulol productivity assessment **(Figure 2C).** Secondary transformation, even with just the reporter, led to the selection of transformants with elevated heterologous patchoulol yields as previously described (Wichmann et al., 2018; Abdallah et al., 2022). However, the greatest improvements were observed when *Pc*PS-mTFP1 constructs were combined with the expression of *Pc*PS-YFP in the same compartment **(Figure 2C).** In the dual compartmentalized context, the cytoplasm-localized 1x*Pc*PS-mTFP1 alone (Plasmid 03) produced 178 fg patchoulol/cell. In contrast, *Pc*PS-mTFP1 + *Pc*PS-mVenus (Plasmids 03 + 08 ‘cytoplasm-cytoplasm’) resulted in an average patchoulol yield of 387 fg patchoulol/cell or 3.2 mg/L with the highest transformant accumulating 424 fg patchoulol/cell, 3.7 mg/L **(Figure 2C)**.

**Figure 2.**
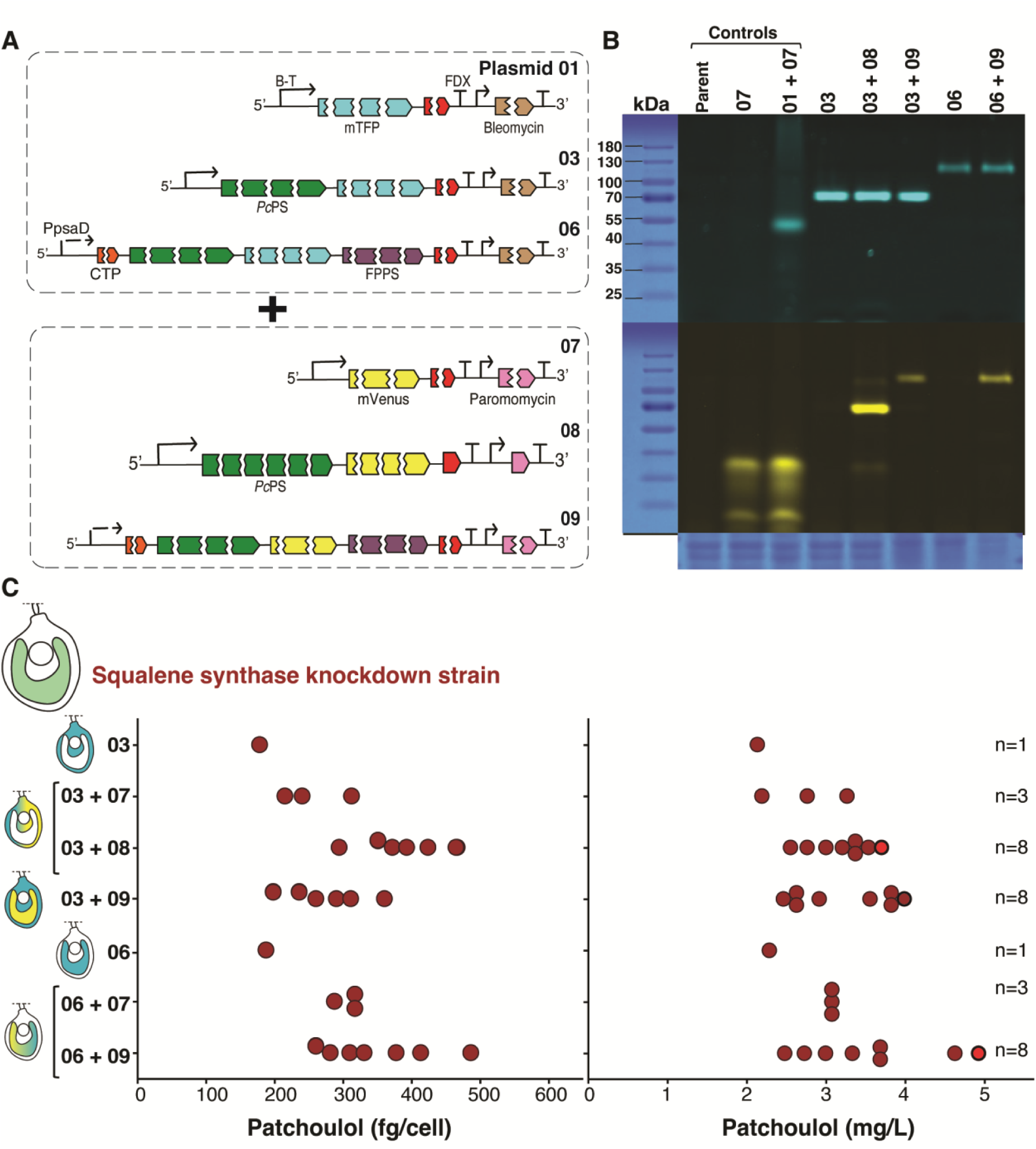
Patchoulol titer differences from enzyme loading in cytoplasm and plastid through double transformations. **A.** plasmid cartoons indicating *Pc*PS expression constructs of plasmids 01, 03, 06-09. Sequences for all plasmids can be found in Supplemental File 01. **B.** SDS PAGE in gel fluorescence of mTFP and mVenus signals for different approximate molecular masses of fusion proteins in representative single and double transformants. **C.** Quantification of patchoulol peaks from GCMS analysis of dodecane taken from cultures after 6 days of cultivation of transformants from the UPN22+SQS k.d. strain. Teal color indicates mTFP fusion expression, yellow, the mVenus fusion expression and the subcellular color indicates where each fusion protein localizes in the algal cell.

Transformation of chloroplast *Pc*PS-FPPS fusion into the cytoplasmic *Pc*PS-mTFP1 strain (Plasmids 03 + 09 ‘cytoplasm-chloroplast’) resulted in an average patchoulol production of 259 fg patchoulol/cell or 3.2 mg/L with highest transformant accumulating 260 fg patchoulol/cell and 3.6 mg/L. We did not identify transformants of this combination with high-level transgene expression, which likely affected overall yields from the cytoplasm-chloroplast combination (**Figure 2B)**. Therefore, these constructs will not be further discussed in this work and were excluded from further analysis **(Figure 2C)**.

However, the most patchoulol was produced by additive transformation of chloroplast-targeted PcPS-FPPS fusions. When a strain with *Pc*PS-mTFP1-FPPS (Plasmid 06) was transformed with *Pc*PS-mVenus-FPPS fusion (Plasmid 09), higher quantities of patchoulol were produced. The chloroplast *Pc*PS-mTFP1-FPPS alone (Plasmid 06) resulted in average patchoulol production of 188 fg /cell or 2.0 mg/L, while the dual transformant (Plasmids 06 + 09 ‘chloroplast-chloroplast’) resulted in an average of 338 fg patchoulol/cell or 4.0 mg/L. The best transformant accumulated up to 413 fg patchoulol/cell or 4.9 mg/L of culture **(Figure 2C).** As the MEP pathway is localized in the chloroplast, the combination of FPPS and the PcPS in this environment likely has an additive effect on patchoulol yields. This is in line with previous findings with bisabolol (Wichmann et al., 2022).

### 3.3. Patchoulol titers with *Pc*PS in cytoplasm or plastid grown on different carbon sources

The interest in utilizing microalgae for bio-production processes stems from their ability to grow using light and CO_2_ as the sole carbon source. However, the alga is also adept at consuming acetic acid as an organic carbon source, which promotes fast growth rates in the light and can even be combined with carbon dioxide for improved growth rates (Stern et al., 2008). Previous research has demonstrated that heterologous terpenoid production in *C. reinhardtii* is variable based on factors such as light, elevated CO_2_ concentrations, and different carbon sources (Lauersen et al., 2016, 2018; Wichmann et al., 2018; De Freitas et al., 2023). The effect of carbon source on patchoulol productivity was investigated in strains with either cytoplasmic or plastid localized *Pc*PS constructs **(Figure 3).** The two transformants were cultivated in 400 mL with 20 mL fluorinated amine underlay to capture patchoulol under three different carbon regimes: 3% CO_2_, acetate, and 3% CO_2_ + acetate **(Supplementary Table 2)**. Cultivations were performed either in constant light or 16 h:8 h light:dark cycles **(Figure 3)**. Growth followed an exponential pattern on acetate, with a stationary phase after three days of cultivation **(Figure 3A).** In CO_2_-supplied cultures without acetate, growth was linear, while cultures provided with CO_2_ + acetate grew exponentially at first followed by a linear later stage. The same trends were observed with lower growth rates when light-dark cycles were used **(Figure 3)**.

**Figure 3.**
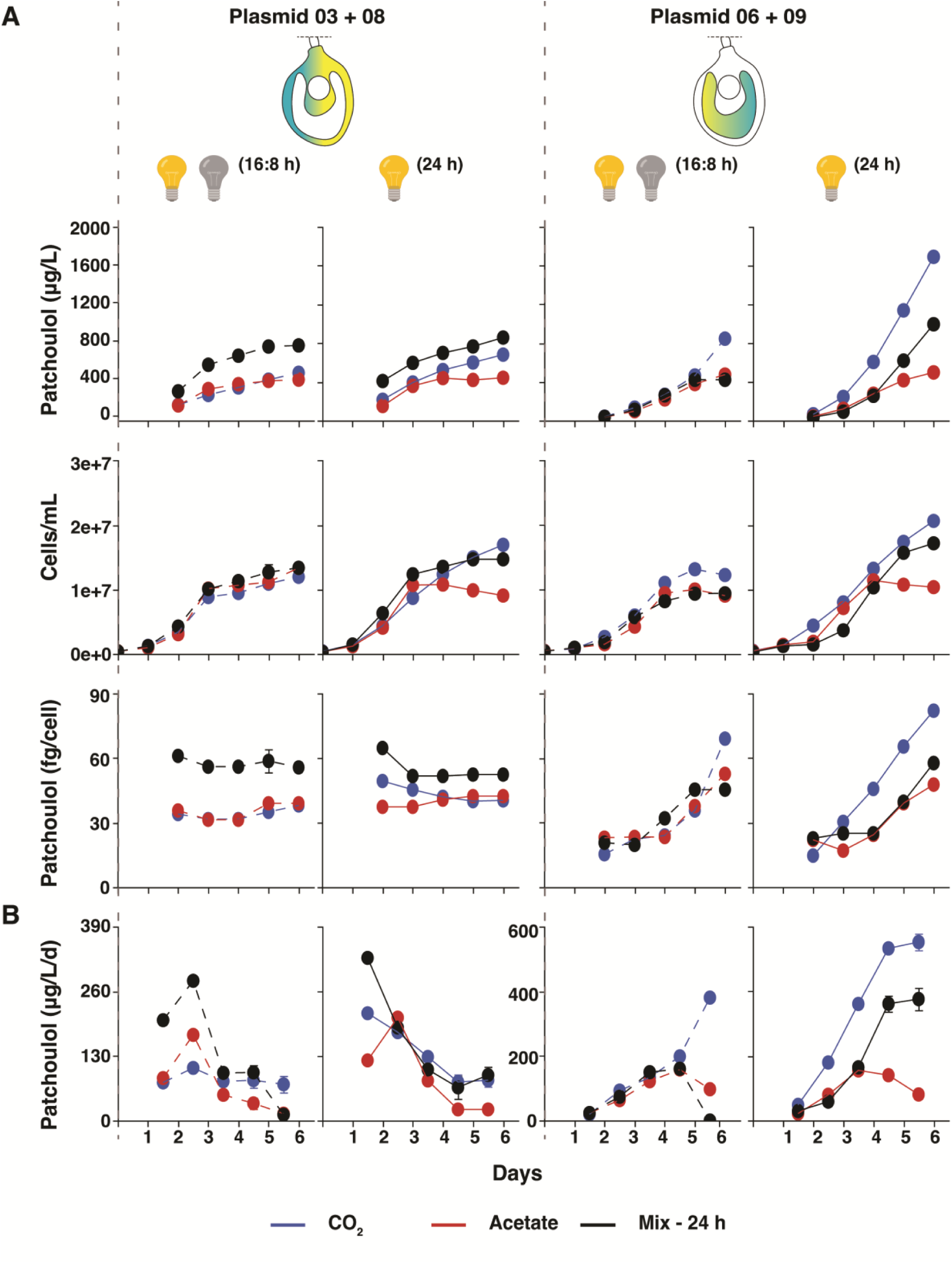
Cultivation of two high producing transformants in photobioreactors and quantification of patchoulol production dynamics. **A**. cultivation of strains was performed in either 16:8 or 24-hour light for two separate strains harboring either plasmids 03+08 or 06+09. Patchoulol from FC3283 underlays was quantified throughout the cultivation with daily sampling, in addition to cell density measurements. **B.** the daily volumetric productivity of patchoulol is shown.

Patchoulol yields differed greatly depending on the carbon source. Over 6 days of cultivation, the transformant with cytoplasm-cytoplasm localized *Pc*PS (Plasmids 03 + 08) produced up to 681 µg patchoulol/L on constant light and 454 µg patchoulol/L in light-dark cycles grown on CO_2_ as sole carbon source. Up to 443 µg patchoulol/L was accumulated in constant light on acetate cultivation, and 383 µg patchoulol/L on a light-dark cycle. In CO_2_ + acetate cultivations, patchoulol production was up to 858 µg/L in constant light and 744 µg/L in light-dark **(Figure 3A).** Across cultivation conditions and carbon sources, volumetric patchoulol production peaked during the early logarithmic growth in the first few days and steadily decreased thereafter **(Figure 3B).**

In the chloroplast-chloroplast *Pc*PS (Plasmids 06 + 09), patchoulol titers were highest in constant light and accumulated up to 1691 µg patchoulol/L when CO_2_ was the sole carbon source, while 846 µg patchoulol/L was produced on light-dark cycles. Up to 497 µg patchoulol/L was accumulated in constant light on acetate, and 475 µg/L was accumulated on a light-dark cycle. In acetate + CO_2_ cultivations, patchoulol was produced up to 995 µg/L in constant light and 422 µg/L on a light-dark cycle **(Figure 3A).** Volumetric yields for all cultivation conditions on constant light and light-dark cycles showed productivity peaks at later cultivation days as opposed to strains with cytoplasm-localized *Pc*PS. An extended patchoulol productivity rate in the later stages of cultivation was observed for the plastid-localized *Pc*PS **(Figure 3B).**

### 3.4. High-density cultivation in CellDEG bioreactors

To assess CO₂-based high-density growth, the two highest patchoulol-producing UPN22-SQS k.d. transformants with different subcellular-localized combinations of *Pc*PS (cytoplasm-cytoplasm #1, chloroplast-chloroplast #5, light red circles of **Figure 2)** were cultivated in CellDEG HD100 bioreactors using nutrient enhanced medium and CO_2_ as the sole carbon source **(Figure 4A).** These bioreactors normally achieve higher cell densities than in 6-well plates or Algem photobioreactors, and potentially higher patchoulol production. After 7 and 14 d of cultivation, patchoulol was quantified by GC-MS/FID **(Figure 4B).** As already observed for the 400 mL photobioreactor experiments, the chloroplast-localized strain outperformed the cytoplasmic strain, accumulating 15 mg patchoulol/L after 14 days, compared to 7 mg/L in the cytoplasmic strain, confirming higher efficiency in chloroplasts **(Figure 4C).** This is currently the highest productivity of patchoulol reported for *C. reinhardtii* (Abdallah et al., 2022; Einhaus et al., 2025; Gutiérrez et al., 2024; Lauersen et al., 2016) and represents a promising step towards heterologous terpene production from CO_2_. Previously, high titers of sclareol diterpenoid were reported in a 2-L version of this system (Einhaus et al., 2022). The sesquiterpene synthases are notoriously catalytically slow enzymes and *Pc*PS is known to produce lower amounts of products than other similar enzymes (Gutiérrez et al., 2025).

**Figure 4.**
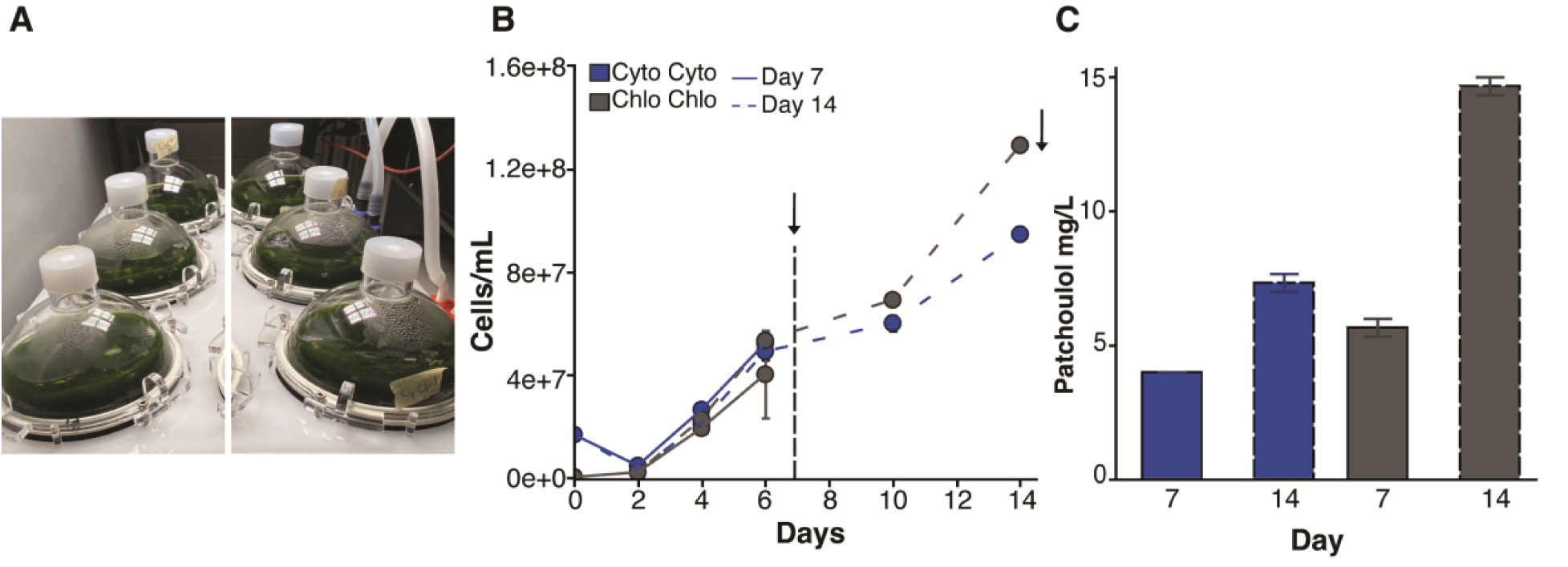
Cultivation of strains producing patchoulol from cytoplasm of chloroplast were compared in high-density photobioreactor cultivation concepts. **A.** photograph of cultures. **B.** quantification of cell densities in photobioreactors from two separate strains harboring either plasmids 03+08 or 06+09. **C.** quantification of patchoulol produced by the algae in their dodecane overlays after 7 and 14 days of cultivation.

## Conclusion

In this study, we aimed to elucidate dynamics in the availability of terpenoid precursors between the cytoplasm and chloroplast of the green alga *C. reinhardtii* by using the sesquiterpene patchoulol as a reporter. Our findings emphasize the importance of subcellular localization in metabolic engineering in *C. reinhardtii* to enhance sesquiterpenoid production. There was no improvement in patchoulol titers within the chloroplast in strains with cytoplasmic SQS knockdown, indicating there is no significant backward flux of FPP to the chloroplast. We demonstrate that farnesyl pyrophosphate pools can be significantly increased in the algal plastid to generate heterologous patchoulol, and it was possible to produce patchoulol production in two cellular compartments simultaneously, but final product yields were highest from chloroplast enzyme localization and under photoautotrophic growth. This approach holds promise for utilizing this microbial host to produce high-value products from waste CO_2_ sources as production was maintained into the later cultivation stages, unlike for cytoplasmic localization. Future engineering efforts can therefore tailor engineering strategies to the intended carbon source.

## Author Contributions

MA, GW, and SO contributed to experimental design and performed experiments. KL was responsible for experimental design, project scope, and funding acquisition. MA, GW, SO, and KL contributed to manuscript preparation.

## Supporting information

Supplemental File 01 - All Plasmid Sequences

Supplemental File 01 - All Data

Supplemental Table 01 - Plasmids used for transformation

Supplemental Table 02 - Photobioreactor Growth Conditions

Supplemental Table 03 - CellDeg cultivation settings profiles

## Acknowledgements

KJL acknowledges baseline research support from King Abdullah University of Science and Technology (KAUST).

