## Supplemental Table 01 - Plasmids used for transformation for "Investigating the dynamics of precursor availability for sesquiterpenoid production in the cytoplasm and plastid of *Chlamydomonas reinhardtii*"

Supplementary Table 1. Genetic constructs used for transformation

| **Plasmid #** | **SSB**  **ID** | **Plasmid name** | **Sel.** | **Rep.** | **Cell loc.** |
| --- | --- | --- | --- | --- | --- |
| **01** | B005 | pOpt3_pAβ_5’btub2_mTFP1_Tfdx1_Ble | B | T | Cyt |
| **02** | B006 | pOpt3_pAβ_5’btub2_mTFP1_ScERG20_Tfdx1_Ble | B | T | Cyt |
| **03** | B019 | pOpt3_pAβ_5’btub2_PcPS_mTFP1_Tfdx1_Ble | B | T | Cyt |
| **04** | B025 | pOpt3_pAβ_5’btub2_PcPS_mTFP1_ScERG20_Tfdx1_Ble | B | T | Cyt |
| **05** | B022 | pOpt3_pPsad_5’psadNoi_CTPpsadi1_1xPcPS_mTFP1_Tfdx1_Ble | B | T | Chlor |
| **06** | B028 | pOpt3_pPsad_5’psadNoi_CTPpsadi1_PcPS_mTFP1_ScERG20_Tfdx1_Ble | B | T | Chlor |
| **07** | P007 | pOpt3_pAβ_5’btub2_mVenus_strepII_Tfdx1_Paro | P | V | Cyt |
| **08** | PA01 | pOpt3_pAβ_5’btub2_PcPS_mVenus_Tfdx1_Paro | P | V | Cyt |
| **09** | P156 | pOpt3_pPsad_5’psadNoi_CTPpsadi1_PcPS_mVenus_Tfdx1_Paro | P | V | Chlor |

Sel. – selection, B – Bleomycin, P – Paromomycin, Rep. – Reporter, T – mTFP1 (cyan),

V – mVenus (yellow), Cell loc. – Subcellular localization, Cyt – cytoplasm, Chlor – Chloroplast
