## Supplemental Table 02 - Photobioreactor Growth Conditions for "Investigating the dynamics of precursor availability for sesquiterpenoid production in the cytoplasm and plastid of *Chlamydomonas reinhardtii*"

Supplementary Table. 2 Photobioreactor growth conditions.

| **Flask#** | **Condition** | **Temperature** | **Light cycle** | **Light intensity μE** | **Media** | **CO_2_ %** |
| --- | --- | --- | --- | --- | --- | --- |
| 1 | CO_2_ only | 23°C | 24h | 325 | T2PhiNO_3_ | 3% |
| 2 | CO_2_ only | 23°C | 16:8h | 325 | T2PhiNO_3_ | 3% |
| 3 | Acetate only | 23°C | 24h | 325 | TA2PhiNO_3_ | - |
| 4 | Acetate only | 23°C | 16:8h | 325 | TA2PhiNO_3_ | - |
| 5 | CO_2_ + Acetate | 23°C | 24h | 325 | TA2PhiNO_3_ | 3% |
| 6 | CO_2_ + Acetate | 23°C | 16:8h | 325 | TA2PhiNO_3_ | 3% |
