## Supplemental Table 03 - CellDeg cultivation settings profiles for "Investigating the dynamics of precursor availability for sesquiterpenoid production in the cytoplasm and plastid of *Chlamydomonas reinhardtii*"

Supplementary Table 3. CellDeg cultivation profiles.

| **Light** | **CO_2_** |
| --- | --- |
| Initial 304 µE | Initial |
| Logarithmic with a doubling time of 36 h until reaching 1000 µE | Linear increase for 12 hrs until 5% |
| Maintained at 1000 µE until day 7 |  |
| Logarithmic with a doubling time of 36 hrs until reaching 2000 µE | Maintained at 5% until end of the experiment |
| Maintained at 2000 µE until Day 14 |  |
